# A Circuit Model of Auditory Cortex

**DOI:** 10.1101/626358

**Authors:** Youngmin Park, Maria N. Geffen

## Abstract

The mammalian sensory cortex is composed of multiple types of inhibitory and excitatory neurons, which form sophisticated microcircuits for processing and transmitting sensory information. Despite rapid progress in understanding the function of distinct neuronal populations, the parameters of connectivity that are required for the function of these microcircuits remain unknown. Recent studies found that two most common inhibitory interneurons, parvalbumin- (PV) and somatostatin-(SST) positive interneurons control sound-evoked responses, temporal adaptation and network dynamics in the auditory cortex (AC). These studies can inform our understanding of parameters for the connectivity of excitatory-inhibitory cortical circuits. Specifically, we asked whether a common microcircuit can account for the disparate effects found in studies by different groups. By starting with a cortical rate model, we find that a simple current-compensating mechanism accounts for the experimental findings from multiple groups. They key mechanisms are two-fold. First, PVs compensate for reduced SST activity when thalamic inputs are strong with less compensation when thalamic inputs are weak. Second, SSTs are generally disinhibited by reduced PV activity regardless of thalamic input strength. These roles are augmented by plastic synapses. These differential roles reproduce the differential effects of PVs and SSTs in stimulus-specific adaptation, forward suppression and tuning-curve adaptation, as well as the influence of PVs on feedforward functional connectivity in the circuit. This circuit exhibits a balance of inhibitory and excitatory currents that persists on stimulation. This approach brings together multiple findings from different laboratories and identifies a circuit that can be used in future studies of upstream and downstream sensory processing.

**Significance Statement:** The mammalian auditory cortex is composed of multiple types of inhibitory and excitatory neurons, which form sophisticated microcircuits for processing and transmitting sensory information. Distinct inhibitory neuron subtypes play distinct functions in auditory processing, but it remains unknown what simple set of underlying mechanisms is responsible for inhibitory cortical function. Here, we built minimal rate and spiking models and identified a specific set of synaptic mechanisms that could best reproduce the broad set of experimental results in the auditory cortex. The simplicity of our model provides an understanding of inhibitory cortical processing at the circuit level, which explains results from different laboratories, and provides for a novel computational framework for future studies of cortical function.

## Introduction

Detecting sudden changes in the acoustic environment and extracting relevant acoustic features from noise are important computations for auditory navigation and scene analysis. The mammalian auditory cortex (AC) is a key region for processing temporally patterned sounds (Ulanovsky, Las, and Nelken 2003). Neurons in AC exhibit adaptation to repeated tones, which may be selective for an overrepresented stimulus, such as in stimulus-specific adaptation, or SSA (Ulanovsky, Las, and Nelken 2003; Natan et al. 2015). They furthermore exhibit forward suppression, in which a preceding stimulus masker tone drives a decrease in responses to the subsequent target tone (Phillips and Hasenstaub 2016; Loebel, Nelken, and Tsodyks 2007). How these computations are carried out by cortical circuits has been subject of extensive research.

The AC is composed of tightly coupled networks of excitatory and inhibitory neurons. Recent studies have identified the differential involvement of two distinct major classes of inhibitory neurons, parvalbumin-positive (PV) and somatostatin-positive (SST) neurons in these temporal paradigms. These neurons differ morphologically and physiologically (Fino, Packer, and Yuste 2013; Isaacson and Scanziani 2011), and recent studies found that they play differential functions in auditory processing. Specifically, SSTs, but not PVs facilitate stimulus-specific adaptation (Natan et al. 2015). PVs and SSTs play distinct roles in adaptation to repeated tones along the frequency response function of the target neuron (Natan, Rao, and Geffen 2017). SSTs and PVs drive bi-directional effects on forward suppression (Phillips, Schreiner, and Hasenstaub 2017). In addition, PVs enhance feedforward connectivity in the auditory cortex (Hamilton et al. 2013). These experimental results can be used to constrain currents and connections in an idealized auditory cortex model consisting of PVs, SSTs and Exc. Here, we tested whether these results can be accounted for by the same set of mechanisms.

In this paper, we build up from a simple dimensionless model consisting of one iso-frequency unit. We transition to a three-unit rate model to understand the mechanisms of inhibitory neural modulation on a gross tonotopy. We then build a detailed spiking model which incorporates the mechanisms discovered in the rate models. The rate models provide a qualitative intuition of the underlying mechanisms, and the spiking models incorporated these mechanisms in an accessible, open-source codebase for future work. We present the results on testing the three-unit rate model and the detailed spiking model in four distinct auditory paradigms.

The model accounted for observed experimental results including the differential role of SSTs and PVs in SSA (Natan et al. 2015), forward suppression (Phillips, Schreiner, and Hasenstaub 2017), tuning-curve adaptation (Natan, Rao, and Geffen 2017), and the effects of PV activation on feedforward functional connectivity (Hamilton et al. 2013). We found that compensating currents between the two types of inhibitory neurons explain experimental findings of differential effects of their modulation on excitatory activity. Furthermore, the model is consistent with existing hypotheses regarding inhibitory and excitatory balance in the cortex. This framework can be used to build and test hypotheses for similar phenomena in other sensory modalities, and studies of upstream or downstream auditory processing in AC.

## Materials and Methods

We first built an augmented version of the Wilson-Cowan model, consisting of one iso-frequency unit of the auditory cortex. The model consisted of one excitatory neural population and two inhibitory neural subpopulations. Importantly, the single iso-frequency unit model served as the template for all other models in this paper. By using the results and parameters from this model, we extended our results to the substantially more complex three-unit rate model and three-unit spiking models. We constrained the parameters using experimental data from the literature.

All code used to generate figures (including model simulations, numerical methods, and analysis methods) are available on GitHub at https://github.com/geffenlab/park_geffen under the MIT open source license.

In figures in this manuscript, we use blue or black lines to depict Exc activity in absence of optogenetic manipulation (called “Control”); magenta for SST activity; cyan solid for PV activity; orange for Exc activity under SST suppression or activation; green for Exc activity under PV suppression or activation.

### Augmented Wilson-Cowan Model

We modeled a single iso-frequency unit as an augmented version of the Wilson-Cowan model (Wilson and Cowan 1972) by including an additional inhibitory subtype. We emphasize that we drew much of our understanding of adaptation throughout this paper using this single iso-frequency unit:

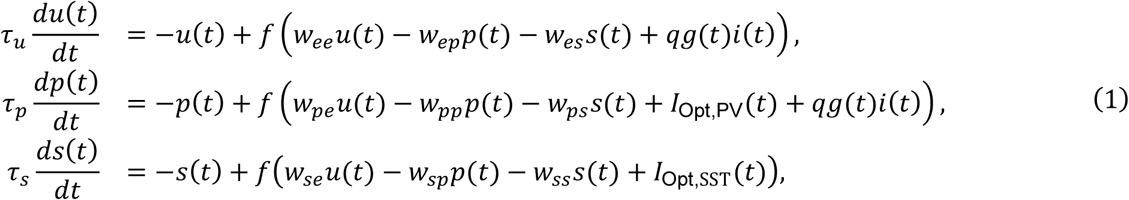

where *u*(*t*), *p*(*t*), and *s*(*t*) represent the normalized firing rate (scaled from 0 to 1) of the excitatory population, PV inhibitory subpopulation, and SST inhibitory subpopulation, respectively (Figure 1. Note that connection strengths in the rate model are given by *w*_*ij*_, and the spiking model are given by conductances *g*_*ij*_, which are distinct from the thalamic depression variable *g*(*t*). We will keep these distinctions consistent throughout the text, so there is no ambiguity). An important caveat of the rate equations is that the rescaled firing rates are a consequence of *dimensionless* equations. All weights, parameters, optogenetic inputs, and thalamic inputs are also dimensionless in the rate model, with the only exception being time.

**Figure 1.**
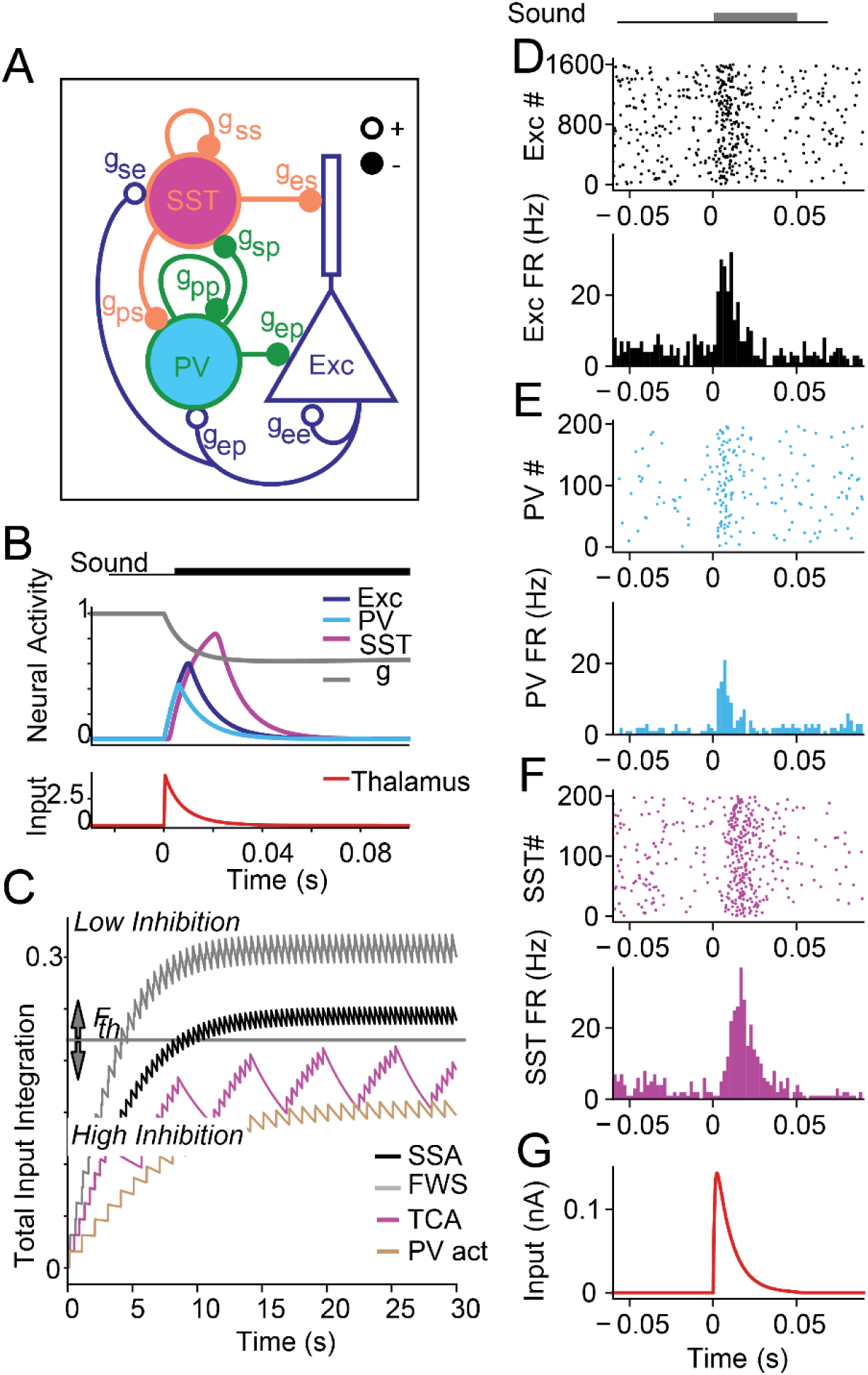
A. Model of the PV-SST-Exc circuit (spiking model, where connection strengths are given by conductances g_ij_. Note that the conductances with subscripts g_ij_ are distinct from g(t), the adaptation variable for the thalamic input. The rate model uses connection strengths w_ij_). B. Input and response profiles for the single-unit rate and spiking model to a 100 ms long tone. Top: Gray: thalamic depression variable g. Blue: excitatory (Exc) neuron activity. Cyan: PV. Magenta: SST. Bottom: Thalamic input (red). C. Responses to stimulus over the first 30 s after sound onset for the different paradigms modeled in the paper. We used a high and a low inhibition mode of synaptic weights to capture the different results. For SSA (black) and forward suppression (FWS, gray), the variable 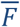 is higher than threshold 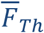, resulting in a set of low inhibition parameters. Paradigms for tuning-curve adaptation (purple) and PV activation (g) asymptote at below-threshold levels, resulting in a set of high inhibition parameters. D-G responses of neurons in the spiking model to a 50ms tone. Top: raster plot; Bottom: Firing rate of Exc (D), PVs (E), SSTs (F), and thalamo-cortical input (G).

The parameters *I*_Opt,PV_(*t*) and *I*_Opt,SST_(*t*) represent the strength of PV and SST activation or inactivation, respectively, and *w*_*ij*_ and *τ*_*i*_ are synaptic weights and time constants, respectively. All time constants are *τ*_*u*_ = *τ*_*p*_ = *τ*_*s*_ = 10ms, roughly in agreement with known data (Tsodyks et al. 1997; Natan et al. 2015). The function *f* is a threshold linear function defined as

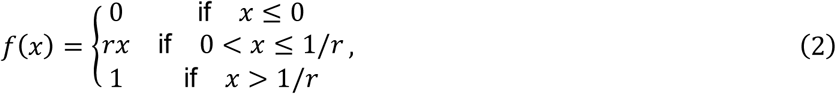

where the function *f* roughly approximates a sigmoid that converges to zero for small or negative inputs and saturates to 1 for large inputs. The parameter *r* = 3 determines the gain of all firing-rate functions and was chosen roughly to be the same as other modeling studies (Yarden and Nelken 2017; Natan et al. 2015). We included a threshold by subtracting a constant *u*_*th*_ from the input, i.e., *f*(*x* − *u*_*th*_) for some input *x*, where *u*_*th*_ is a positive number (typically in the range from 0 to 1). In all rate model simulations, we chose Exc, PV, and SST thresholds to be *u*_*th*_ = 0.7, *p*_*th*_ = 1, and *s*_*th*_ = 1, respectively. The thresholds indicate the minimum activity required for a neural population to affect postsynaptic neural populations. Because the thresholds are greater than zero, sub-threshold activity does not affect the dynamics of the network.

The input function *i*(*t*) consists of blocks of inputs with stimulus duration and interval based on the experimental paradigm. We list the stimulus duration and stimulus interval for each paradigm in Table 2 and detail the paradigm in the text and figures where appropriate. When an auditory input arrives into the Exc and PV populations, the default temporal profile is taken to have an instantaneous rise with amplitude *q* and exponential decay (Figure 1B, bottom red curve) with time constant *τ*_*q*_ = 10ms, which roughly agrees with known values (Rankin, Sussman, and Rinzel 2015). The instantaneous rise and exponential decay were chosen for simplicity. The input *i*(*t*) is further modulated by a slow synaptic depression term *g* satisfying the standard model of synaptic depression

**Table 1:** Parameter values of rate models across paradigms. In the optogenetic parameters, numbers without parentheses show optogenetic strengths for use in model reproductions and numbers in parentheses show optogenetic strengths for use in model predictions. Positive optogenetic numbers correspond to activation and negative numbers correspond to inactivation.

| Parameter | SSA (Simple) | SSA | FWS | TCA | PV Act. | Notes |
| --- | --- | --- | --- | --- | --- | --- |
| $q$ | 5 | 5 | 1.3 | 5 | 5 | Fitted, dimensionless, rate |
| $I_{\text{Opt,PV}}(t)$ | -2 | -4 (0.5) | -0.5 (0.25) | -0.5 (1.2) | 2 (2) | Fitted, dimensionless, rate |
| $I_{\text{Opt,SST}}(t)$ | -1 | -2 (1.2) | -0.5 (0.1) | -1 (0.1) | - | Fitted, dimensionless, rate |
| q | 5 | 5 | 1.3 | 5 | 5 | Fitted, dimensionless,<br>spiking |
| $I_{Opt,PV}(t)$ | - | -0.2nA | 0.1nA | 1nA | 0.5nA | Fitted, dimensionless,<br>spiking |
| $I_{Opt,SST}(t)$ | - | -1nA | 2nA | 1nA | - | Fitted, dimensionless,<br>spiking |

**Table 2:** Auditory paradigm parameters. SSA parameters from (Natan et al. 2015). Forward suppression (FWS) parameters from (Brosch and Schreiner 1997; Phillips, Schreiner, and Hasenstaub 2017). Tuning-curve adaptation (TCA) parameters from (Natan, Rao, and Geffen 2017). PV activation parameters from (Hamilton et al. 2013). Optogenetic inhibition was performed 100ms before tone onset and 100ms after tone offset in SSA and FWS. In TCA and PV activation, optogenetic inhibition was turned on at the beginning of the experiment and sustained through the trial. For optogenetic parameters, see Table 2.

|  | SSA | FWS | TCA | PV Act. |
| --- | --- | --- | --- | --- |
| Stimulus duration | 100ms | 50ms | 100ms | 50ms |
| Inter-stimulus interval | 300ms | 20ms | 300ms | - |
| Inter-trial interval | - | 380ms | 2400ms | 950ms |
| Stimuli per trial | - | 2 | 8 | 1 |

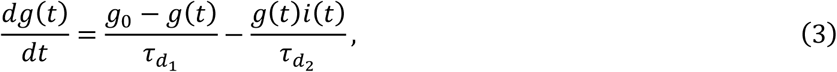

where the time constants are 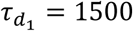 ms for replenishment and 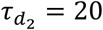ms for depletion (chosen close to reported values (Abbott et al. 1997; Wehr and Zador 2005; Tsodyks et al. 1997; Natan et al. 2015)). The synaptic depression variable *g* begins at a baseline value of *g*_0_ = 1 and when *i*(*t*) > 0, i.e., when an input arrives into the Exc or PV populations, *g*(*t*) decreases on the timescale determined by 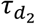. Because *g*(*t*) multiplies the input *i*(*t*) to *u*(*t*) and *p*(*t*) in Equation (1), *g*(*t*) serves to modulate the strength of auditory inputs to A1. In the absence of auditory input, *g*(*t*) recovers slowly on the order of seconds determined by 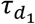.

We based the proportional strengths of connections in the single-unit model based on existing studies on AC (Pfeffer et al. 2013). The within-unit connectivity is equivalently represented by the matrix,

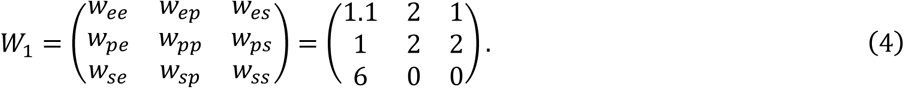

All synaptic weights *w*_*ij*_ in the single-unit rate model are constant (Mill et al. 2011), with synaptic depression appearing in the feedforward thalamic inputs (Lee and Sherman 2008). The inhibitory synaptic weights were roughly chosen to agree with known connection types and connection strengths (Womelsdorf et al. 2014; Pfeffer et al. 2013), and the excitatory connections were tuned as free parameters. The constant synapses allowed us to fully understand the model dynamics before transitioning to the more complex three-unit model with depressing and facilitating synapses.

### Three-unit Rate Model

Using the single-unit rate model as a template, we arranged copies into three units with lateral cortical and thalamic connections (Figure 2A. Lateral inhibitory connections are hidden in Figure 2 for clarity). This arrangement endowed our model with a gross tonotopy, which we used to explore spectrally and temporally complex auditory inputs. While the three-unit rate model appears to be substantially more complex, the parameters were strongly constrained by the single-unit rate model. In particular, we aimed for each unit of the three-unit model to mimic the excitatory and inhibitory currents of the single-unit rate model. We found that maintaining currents explained many of the known optogenetic experiments in the literature. Before turning to the spiking model, we briefly describe technical details of the parameter values and functions. Each unit behaves according to the equations

**Figure 2.**
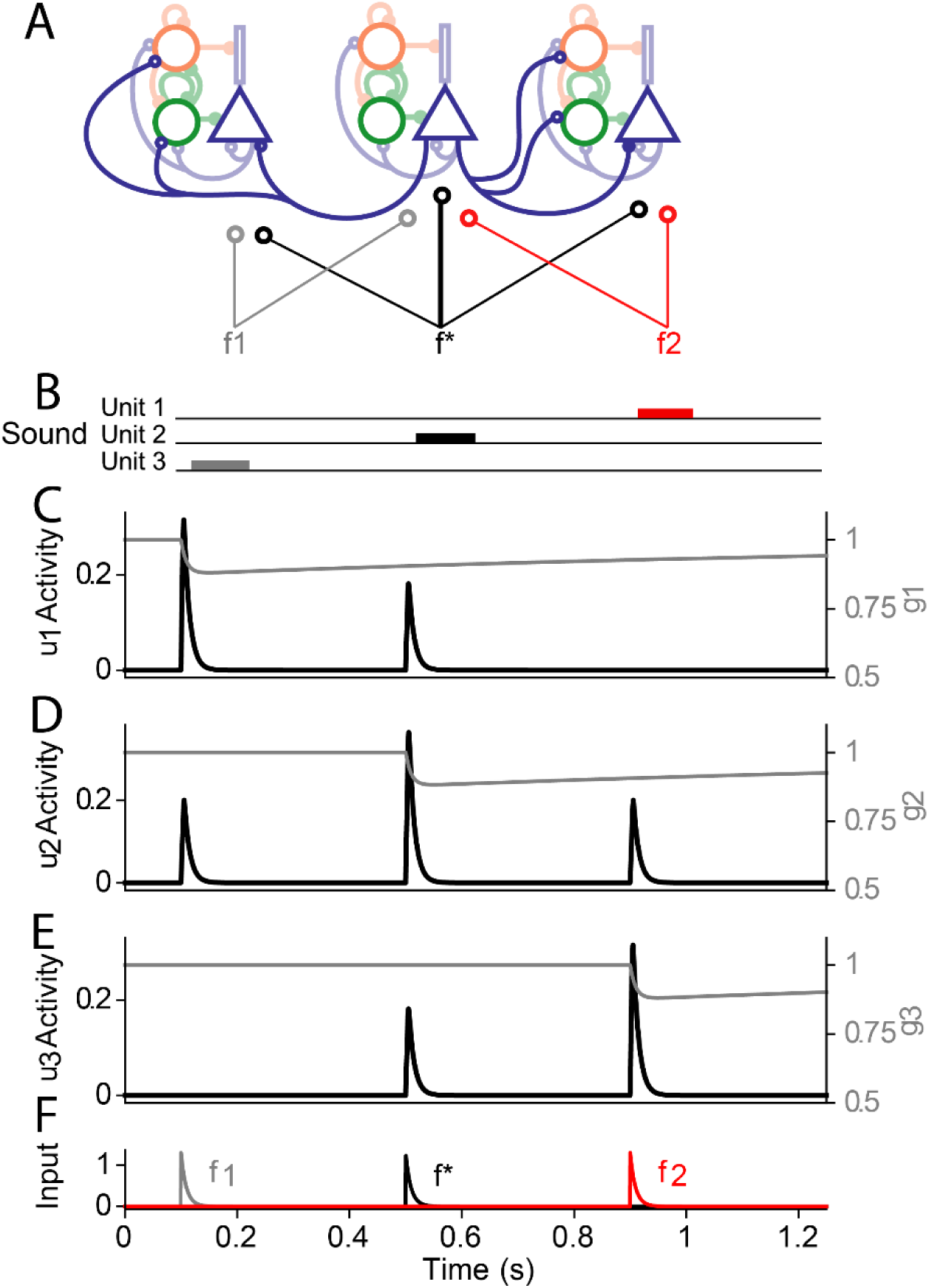
Input and response profiles of the three-unit model. A: The three-unit rate model of the auditory cortex, with three preferred frequencies, f_1_, f^*^, and f_2_ (the spiking model follows the same motif). Non-excitatory lateral connections have been hidden for clarity. B: 50ms auditory inputs are applied at each frequency in sequence. C—E: Black traces show the excitatory cortical response of the first (u_1_), second (u_2_), and third (u_3_) rate units, respectively. Gray traces show the slow synaptic depression. F: The traces of the thalamic inputs: f_1_ (gray), f^*^ (black), and f_2_ (red). Each iso-frequency unit contains lateral excitatory connections where the Exc population of a given unit synapses laterally onto the neighboring Exc, PVs, and SSTs.

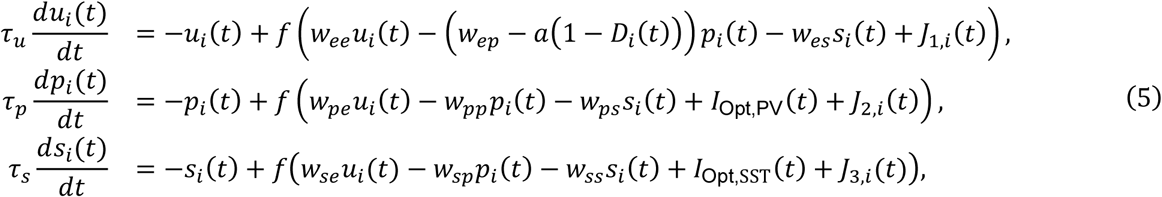

where,

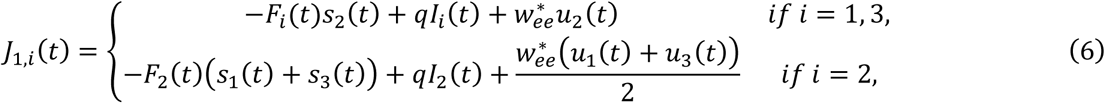

and

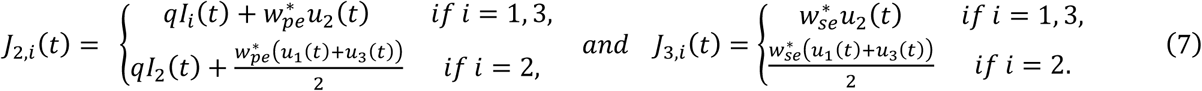

The functions *I*_*i*_(*t*) are defined as *I*_*k*_(*t*) = *g*_*k*_(*t*)*i*_*k*_(*t*) + *g*_2_(*t*)*i*_2_(*t*)*α*, for *k* = 1,3, and *I*_2_(*t*) = (*g*_1_(*t*)*i*_1_(*t*) + *g*_3_(*t*)*i*_3_(*t*))*α* + *g*_2_(*t*)*i*_2_(*t*). We remark that any lower-case letter *i* that has a subscript and is a function of time, e.g., *i*_1_(*t*), *i*_2_(*t*), *i*_3_(*t*), represent thalamic inputs. The time-dependent notation for these functions will always be distinct from the index *i*. Note that each set of equations are almost identical to the single-unit case, but with the addition of lateral terms along with facilitating and depressing terms *F*_*i*_(*t*) and *D*_*i*_(*t*). The lateral terms are between immediate neighbors and include lateral SST to Exc (facilitating), Exc to Exc, Exc to PV. The facilitating terms *F*_*i*_(*t*) increase from 0 to nonzero values as unit *i* receives inputs, and the depressing terms *D*_*i*_(*t*) decrease from 1 to lower values as unit *i* receives inputs. Rather than adjusting weights integrated prior to facilitating synapses, we controlled the strength of facilitation by changing the time constants in Equation (8).

We chose *α* = 0.65, i.e., 65% of the thalamic inputs to the left or right units reach the center unit. Likewise, 65% of thalamic inputs to the center unit reach the left and right units. The function *f* is threshold linear (Equation 2). The functions *I*_*k*_ (*t*) are time-dependent inputs with the strongest preference for unit *k*, and the profiles of *i*_1_(*t*), *i*_2_(*t*), and *i*_3_(*t*) are shown in Figure 2F (these profiles are the same as the profile in the single-unit model, Figure 1B, bottom). Parameters *a, b* control the strength of depression and facilitation and are chosen to be *a* = 0.5, *b* = 2. The parameter *q* controls the strength of all inputs. Each input *I*_*j*_(*t*) is modulated by corresponding depression variables *g*_*k*_(*t*), where each *g*_*k*_(*t*) satisfies Equation 3 independently. The parameters *τ*_*i*_ are membrane time constants and chosen the same as the single-unit model, *τ*_*u*_ = *τ*_*p*_ = *τ*_*s*_ = 10ms (Tsodyks et al. 1997; Natan et al. 2015). The parameters *w*_*ij*_ are within-unit synaptic weights chosen according to Equation 4, while the parameters 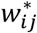are lateral (between unit) synaptic weights. We chose 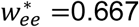, 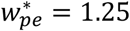, and 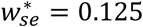 to reflect the generally weaker lateral synaptic strengths in auditory cortex relative to the within-unit connections (Levy and Reyes 2012).

We added facilitating terms *F*_*i*_(*t*) in the Exc to SST synapses, and depressing terms *D*_*i*_(*t*) in the PV to Exc synapses (Beierlein, Gibson, and Connors 2003). The parameters *a* and *b* control the degree of depression and facilitation, respectively, where we chose *a* = 0.5, *b* = 2 (these values were not taken from the literature). The depressing parameter *a* was chosen so that the term (*w*_*ep*_ − *aD*_*i*_(*t*)) did not change sign across experimental paradigms. The facilitating variables *F*_*i*_(*t*) satisfy

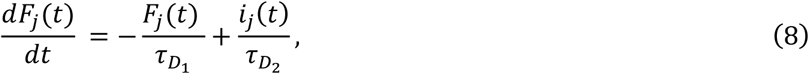

where 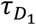 and 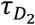 are as in Equation 3. The reuse of the depression time constants 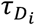 are an intentional and simplifying choice. By using the same time scales, we were able to use the inputs *i*_*j*_(*t*) as a proxy for the excitatory activity *u*_*j*_(*t*) and simulate the facilitation variable in terms of the depressing synaptic variable as *F*_*j*_(*t*) = 1 − *g*_*j*_(*t*). Similarly, the depression variables *D*_*i*_(*t*) satisfy

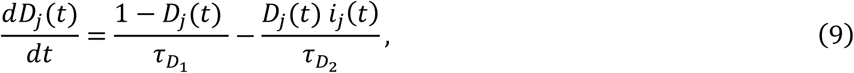

and used the thalamic input as a proxy for excitatory activity to simulate the depression variable as *D*_*i*_(*t*) = *g*_*i*_(*t*). All depressing and facilitating timescales were chosen close to reported values (Abbott et al. 1997; Tsodyks et al. 1997; Wehr and Zador 2005).

The activity of the model is shown in Figure 2, in response to three successive auditory stimuli are applied in order of the frequencies *f*_1_, *f*^*^, and *f*_2_, stimulating the left, center, and right units, respectively (Figure 2C-E). The center unit (*u*_2_(*t*)) responded equally well to frequencies *f*_1_ and *f*_2_ (Figure 2D), a necessary response for SSA paradigms. For simplicity, activation of an adjacent unit did not affect the thalamic variable, i.e., *g*_1_(*t*) was left unaffected by *i*_2_(*t*), *g*_2_(*t*) was left unaffected by *i*_1_(*t*) and *i*_3_(*t*), and *g*_3_(*t*) was left unaffected by *i*_2_(*t*). We assumed that the frequency difference between *f*_1_ and *f*_2_ was great enough that auditory inputs at *f*_1_ (*f*_2_) did not affect units responsive to *f*_2_ (*f*_1_).

When matching the model outputs to experiments in the literature, we found that there were seemingly two different parameter sets that explained all phenomena and we were unable to tweak the rate model to unify the parameters. We unified the disjoint parameter sets by incorporating paradigm-dependent baseline states in the three-unit rate model. The model parameters switch between weak and strong baseline inhibition, where weak inhibition corresponds to high thalamic activity (which corresponds to the parameters in Equation (4)), and strong inhibition corresponds to relatively low thalamic activity (which corresponds to the parameters in Equation (11)). This idea was implemented using an additional facilitating variable,

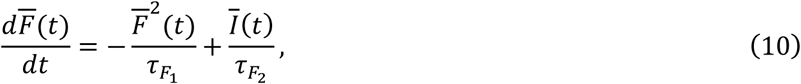

where *Ī* (*t*) represents all thalamic inputs of any frequency, and the parameters are chosen to be 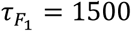, and 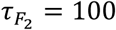. As we applied auditory inputs from particular experiments into the model, the function 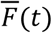 grew as a function of the excitatory input function *I*(*t*), and eventually saturated to different mean values based on the stimulus duration, stimulus interval, and inter-trial interval. Incidentally, we found that the facilitating variable 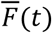 saturated to greater values for forward suppression and SSA, and saturated to relatively lower values for PV activation and tuning-curve adaptation. The two putative disjoint parameter sets correspond exactly to these relatively high and low values of the variable 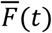. A simulation of Equation 10 is shown in Figure 1C for the various auditory paradigms, with a horizontal line shown where we chose to separate the saturation values.

If 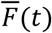 is above the threshold, which we chose to be *F*_*th*_ = 0.22, the system exhibits weak baseline inhibition, and the synapses take baseline values as shown in Equation 4. On the other hand, if 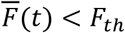, the synapses take the strong baseline inhibitory values

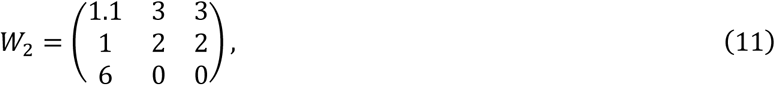

and the SST activity threshold, *s*_*th*_ = 1, decreases to *s*_*th*_ = 0. In our original model, we chose a smooth transition between these parameter sets, i.e., *w*_*ep*_(*t*) = 2*h*(*F*(*t*)) + 3(1 − *h*(*F*(*t*))), 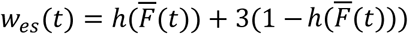, and 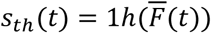 where

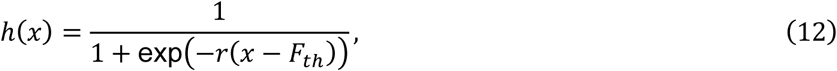

and *r*, the gain of the sigmoid *h* was chosen to be steep, e.g., *r* = 25. However, for simplicity, we replaced *h* with a Heaviside function and assumed that the system already reached either the weak baseline inhibition *W*_1_ (Equation 4), or the strong baseline inhibition *W*_2_ (Equation 11) based on the given experimental paradigm.

For each paradigm (with paradigm parameters shown in Table 2, we simulated Equation 10 and found that SSA and forward suppression belonged to the weak inhibitory regime (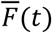 integrated to values above threshold *F*_*th*_), whereas tuning-curve adaptation and PV activation belonged to the strong inhibition regime (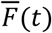 integrated to values below threshold *F*_*th*_). All rate models were simulated using the dynamical systems software XPP (Ermentrout 2002), called using PyXPPCALL and visualized using Python (Rossum 1995). The equations were well-behaved enough that a very coarse time step of dt=0.1 was sufficient for our purposes. We used the default integration method in XPP, the fourth-order Runge-Kutta.

### Goodness of Fit and Related Metrics

We compared the results obtained with the model as result of optogenetic perturbations to published data qualitatively, by testing whether the model reproduces the increase/decrease of activity that was reported. The limit of large numbers yields theoretically tractable equations such as the Wilson-Cowan models we use in this paper. However, finite size effects may contribute to additional issues that the large-number limit does not address. How do we know that the observed changes would hold consistently with the inclusion of finite and noisy neural populations? To address this question, we built a spiking model constrained by the rate models above. Because the spiking model is derived directly from the simpler rate models, we generally expect that the spiking models will reproduce the rate models results for sufficiently large neural population s, however, we are able to establish that the results are consistent in the presence of noise. Moreover, we built the rate model using Python and brian2, which are both extensively documented and accessible open-source packages. Finally, we include the spiking model as a means for researchers to directly fit parameters from experimental data if needed. The rate model provides strong qualitative intuition, but does not explicitly account for single-cell interactions. Thus, the spiking model provides an additional, detailed framework for others to modify and extend beyond what the rate model may provide.

### Spiking Neuron Dynamics

We use the single-unit rate model as a template for the spiking model, constraining the parameters while preserving the pattern of excitatory and inhibitory currents. All inhibitory neurons consisted of a single somatic compartment, while the excitatory neurons were modeled as two-compartment, “ball-and-stick” models. For each excitatory and inhibitory neuron, we modeled the somatic (ball) compartment as an adaptive exponential integrate-and-fire neuron (Brette and Gerstner 2005; Litwin-Kumar, Rosenbaum, and Doiron 2016):

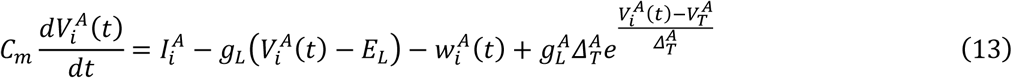

where the transmembrane currents are 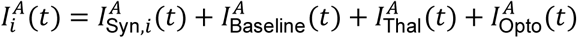, where

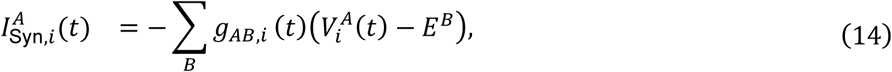

and the sum ∑_*B*_ in the synaptic current iterates over the presynaptic neurons, *B* ∈ {*e, p, s*}. If the presynaptic neuron *B* is excitatory or inhibitory, then *E*^*B*^ = 0mV or −67mV, respectively. If a synaptic connection existed from PV to Exc, we included a depression variable, *D*(*t*), satisfying Equation 9, with 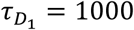, and 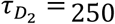:

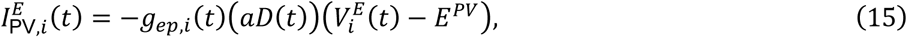

where *a* = 1.7. As in the rate model, the parameter *a* was chosen such that the sign of 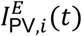 did not change. The additional depression term was necessary to incorporate depression effects that operate well beyond the timescale of inhibitory conductances (Wehr and Zador 2005).

We added transmembrane noise in the form of a white noise process with zero mean and a standard deviation of 20mV to simulate intrinsic and extrinsic current fluctuations. All fixed parameters for each neuron type are shown in Table 3. The parameters that we varied manually were entirely contained in the time-dependent functions 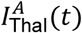 (thalamic inputs) and 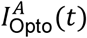 (optogenetic parameters). The thalamic input profile, 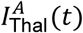, is determined by

**Table 3:** Parameter values of spiking neurons.

|  | Exc | Dend | PV | SST |
| --- | --- | --- | --- | --- |
| $C_m$ (pF) | 180 | 180 | 80 | 80 |
| $E_L$ (mV) | -60 | -60 | -60 | -60 |
| $g_L$ (nS) | 6.25 | 6.25 | 5 | 5 |
| $\Delta_T$ (mV) | 1 | - | 0.25 | 1 |
| $V_T$ (mV) | -40 | - | -40 | -45 |
| $V_{\text{reset}}$ (mV) | -60 | - | -60 | -60 |
| $g_{sd}$ (nS) | 18.75 | 18.75 | - | - |
| $I_{\text{baseline}}$ (nA) | 0.35 | - | 0.05 | 0.025 |

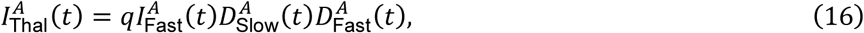

where

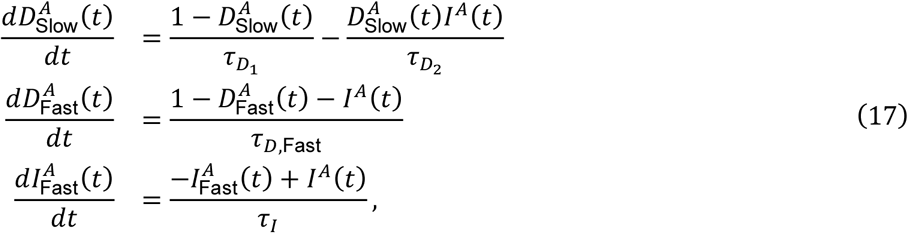

where *τ*_*I*_ = 1ms, *τ*_*D*,Fast_ = 10ms, 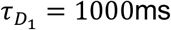, and 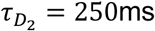. The functions *I*^*A*^ (*t*) (distinct from 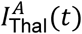 are square wave functions that are active for the duration of the auditory stimulus. Just as in the rate model, the thalamic input function 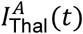 only appears in Exc and PV neurons. The profile of the thalamic input is shown in Figure 1G. The optogenetic term, 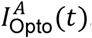, only appears in the PV and SST equations.

Following a presynaptic spike from neuron *j*, the postsynaptic effect on neuron *i* appears as an instantaneous spike in the postsynaptic conductance *g*_*ij*_(*t*) → *g*_*ij*_(*t*) + *g*_*ij*,max_/*n*_X_, where *g*_*ij*,max_ is given by Equation 18, and X stands for the presynaptic neuron type (Exc, PV, or SST). The magnitude of the conductances were chosen to have the same proportion as reported values (Pfeffer et al. 2013), with the same type of connectivity structure as in the rate model

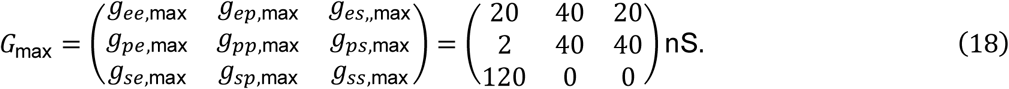

In the absence of presynaptic spikes, the conductances *g*_*ij*_ decay exponentially to zero:

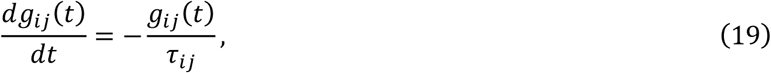

where *τ*_*ij*_ = 1ms for all synapses except for the time constants from excitatory to PVs, *τ*_*pe*_ = 25ms, and excitatory to SSTs, *τ*_*se*_ = 15ms (Markram, Wang, and Tsodyks 1998). In the spiking model, we switched to the weak inhibitory regime by decreasing the inhibitory inputs into Exc from *g*_*ep*,max_ = 40 and *g*_*es*,max_ = 20 to *g*_*ep*,max_ = 38 and *g*_*es*,max_ = 19.

For excitatory neurons (*A* = *e*), the transmembrane currents are 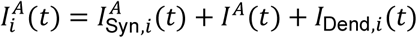,

where

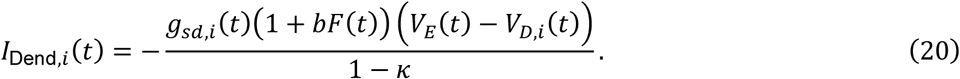

The term *F*(*t*) is a dimensionless slow timescale facilitation variable that depends on the thalamic drive, and satisfies Equation 8 (just as in depression, the additional slow timescale allows the model to operate on multiple timescales (Wehr and Zador 2005)). The parameter *b* = 3 modulates the facilitation strength, and 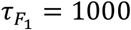, and 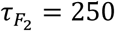. For simplicity, we allowed *F*_*i*_(*t*) to vary continuously over time. The variable *w* represents spike-frequency adaptation and obeys

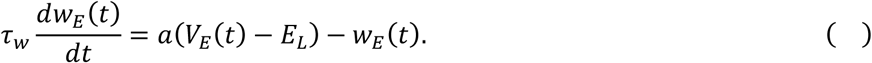

The dynamics of the dendritic (stick) compartment obey

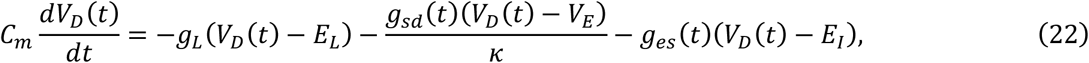

where the parameter *κ* = 0.3 is the ratio of somatic to total surface area (Litwin-Kumar, Rosenbaum, and Doiron 2016).

For PV and SST interneurons, the equations are the same as Exc except that there is no dendritic component. Parameters differ marginally between PV and SST neurons (see Table 3). SSTs, unlike PVs, have no incoming synaptic connections from the thalamus and only receive excitatory inputs from Exc. PVs and SSTs include the optogenetic stimulus term 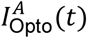, and as mentioned above, only Exc and PVs contain the thalamic input term 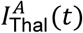. These connections reflect the choices made in the rate model.

### Three-unit Spiking Model

We introduced the gross tonotopy into the spiking model by copying the single unit spiking model into three units with lateral excitatory connections (Figure 2A). As in the 3-unit rate model, the thalamic inputs of the 3-unit spiking model have weaker lateral connections. For tone responses at frequency *f*_1_ and *f*_2_, the center unit receives an input of amplitude proportional to 0.85 that of the left and right units, respectively.

The spiking model contains 1600 Exc, 200 PVs, and 200 SSTs. For connection probabilities within units, we chose *E ← E* connections to have probability *p*^*EE*^ = 0.1 and all other probabilities to be the same, *p*^*EE*^ = *p*^*ES*^ = *p*^*PE*^ = *p*^*PP*^ = *p*^*PS*^ = *p*^*SE*^ = 0.6. For lateral connection probabilities, we chose *p* = 0.1. The spiking model was constructed using Brian2 (Stimberg, Brette, and Goodman 2019).

## Results

### Differential effects of interneuron suppression in stimulus-specific adaptation

Almost all neurons (95%) in AC exhibit stimulus-specific adaptation, a phenomenon in which neurons reduce their response selectively to the inputs that is presented frequently in the stimulus (standard tone), while preserving the initial strong response to the less frequent input (deviant, or oddball tone) (Ulanovsky, Las, and Nelken 2003). Previous studies found that following a presentation of the deviant tone, the excitatory neurons adapt over successive presentations of the standard (von der Behrens et al. 2009; Ulanovsky, Las, and Nelken 2003). Similar adaptation was observed in the songbird (Bell, Phan, and Vicario 2015; Lu and Vicario 2017). This phenomenon was largely attributed to feedforward thalamo-cortical depressing synapses (Mill et al. 2012; 2011), but such models could not account for the full range of the effects that were observed (Yarden and Nelken 2017). A recent study found that inhibitory neurons exhibit differential control over SSA (Natan et al. 2015). Suppressing SSTs resulted in disinhibition of the excitatory neurons, such that disinhibition increased with successive presentations of the standard tone, but not the deviant. By contrast, PV inhibition drove equal amount of disinhibition of excitatory neurons in response to both the deviant and the standard. These results suggest that SST inhibition controls adaptation level of excitatory neurons.

In order to understand the roles of inhibitory interneurons in modulating SSA responses, we tested whether the single-unit model could reproduce the differential effects of suppressing PVs and SSTs in temporal adaptation. This paradigm involved the application of 8 successive tones to the single iso-frequency circuit with constant synapses and depressing thalamic inputs (Figure 3, Equation 1). We then tuned the parameters of the model in order to achieve the qualitative results for SSA (Natan et al. 2015). We simulated optogenetic suppression of inhibitory neurons by defining the functions *I*_Opt,PV_(*t*) and *I*_Opt,SST_(*t*) to turn on 100ms before tone onset and turn off 100ms after tone onset, thereby inhibiting the activity of PV or SST neurons as performed in the experimental paradigms. The degree of PV and SST inhibition was chosen as a free parameter, and in the case of this single-unit rate model, had the dimensionless values of *I*_Opt,PV_(*t*) = −2 during PV suppression and *I*_Opt,SST_(*t*) = −1 during SST suppression. Through this suppression of inhibition, we sought to reproduce the constant disinhibition from PV inactivation, and increasing disinhibition from SST inactivation. Once the model reproduced this qualitative result, we discovered that the key mechanisms involve the temporal structure of the responses: PVs exhibit a temporally fast tone-evoked response and peak earlier than Exc and SSTs, while SSTs exhibit a temporally delayed and broad tone-evoked response (Figure 1B, Figure 3B top left plot), which both agreed with earlier studies (Natan et al. 2015). The SST delay is not hard-coded, but the result of SSTs receiving indirect thalamic excitation through the Exc population (Pfeffer et al. 2013).

**Figure 3.**
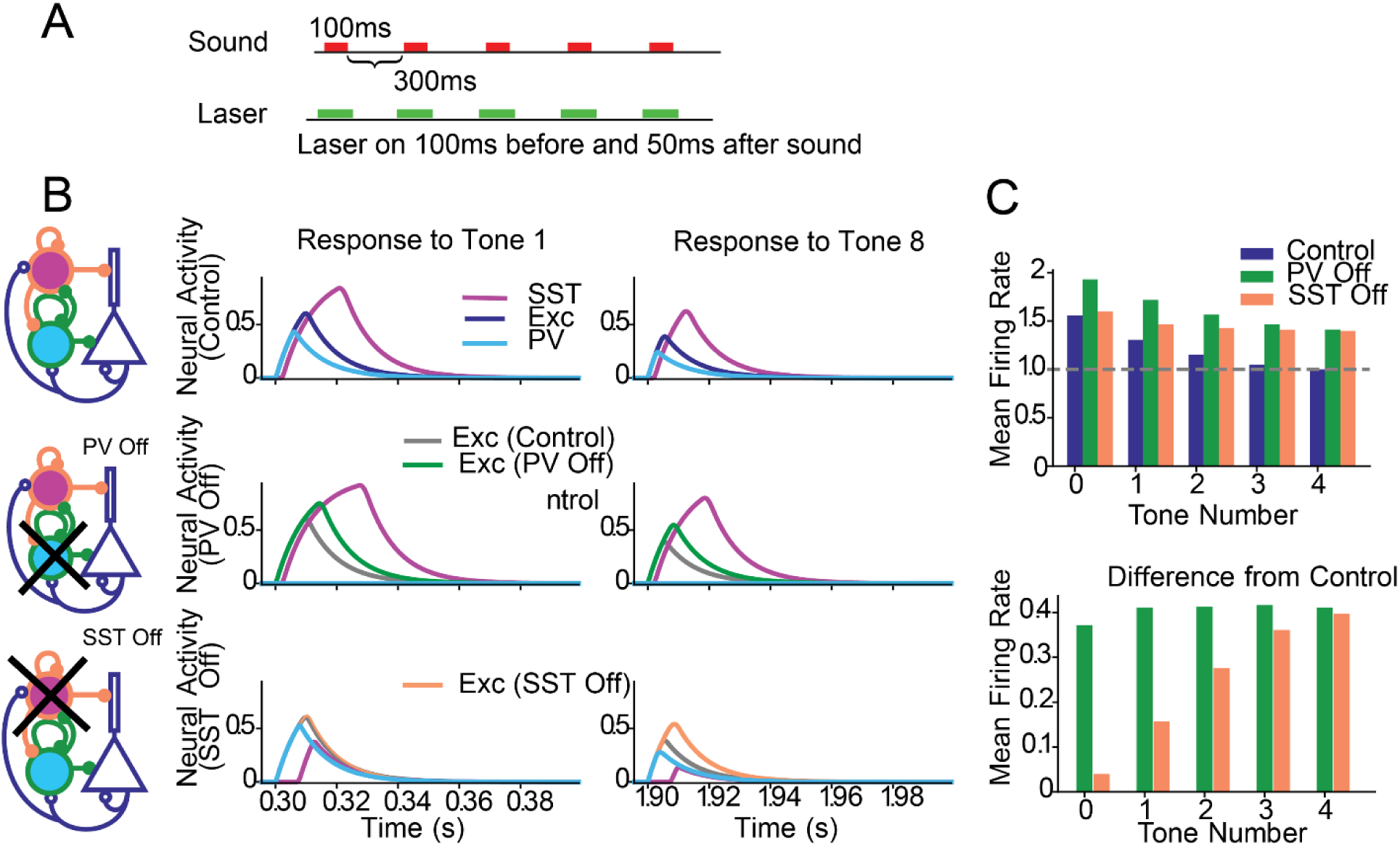
The effect of optogenetic manipulations on adaptation to repeated tones. A. Stimulus of repeated tones, with or without concurrent laser stimulation. B. Left column: Circuit diagram specifying the inactivation of populations. Top row: no stimulation. Responses of Exc (blue), PV (cyan) and SST (magenta) populations to the first (middle column) and last (right column) tones. Middle row: responses during PV suppression. Green: Responses of Exc under PV suppression, gray: responses of Exc in the control condition. Bottom row: responses during SST suppression. Orange: Responses of Exc under SST suppression, gray: responses of Exc in the control condition. C. Top: Mean response of the excitatory population to the repeated tones. Bottom: difference in excitatory responses with and without stimulation. No stimulation (blue), PV suppression (green); SST suppression (orange).

The simplicity of the model allowed us to understand the mechanism underlying these changes. One important observation is the faster temporal activation of PVs relative to SSTs. With this property, excitatory activity is immediately affected by changes to PV activity and less by changes to SST activity. With PV suppression (Figure 3C middle row), PVs are reduced in activity, leading to greater Exc activity. The SSTs were indeed disinhibited due to a lack of inhibition from PVs and the greater Exc activity, but the inhibition from SST to Exc was not strong enough to compensate for changes in PV activity. Thus, the Exc population received an overall decrease in inhibitory current across all successive tones, resulting in constant disinhibition prior to and following adaptation (Figure 3D green). This result suggests that the temporally faster PV inhibition of Exc neurons is necessary for modulating excitatory activity, and only a few milliseconds of earlier PV activity results in a substantially stronger inhibitory effect relative to SSTs.

With SST inactivation, Exc activity at the first tone was virtually unaffected because the reduced SST activity resulted in PV disinhibition, and the increased PV activity resulted in no net change to the total inhibitory current entering the Exc population (Figure 3C, bottom left). PVs compensated for the reduced SST activity. Following adaptation, the overall reduced excitatory activity in both thalamus and Exc resulted in reduced PV activity and a net loss of inhibitory current in Exc. The Exc population was therefore disinhibited at the last tone (Figure 3C, bottom right). Combined, these effects drove an increase in disinhibition over successive tones (Figure 3D, orange).These simple mechanisms of disinhibition and compensation can therefore explain the complementary roles of inhibitory interneurons in shaping cortical activity.

To understand the model dynamics in the presence of inputs at multiple frequencies in the oddball paradigm, we extended the model to a rate and spiking model with three iso-frequency units, in which each microcircuit received inputs of specific preferred frequencies. The three-unit circuitry was based on the single-unit model and the free parameters tuned to reproduce the inhibitory and excitatory currents. For example, an auditory input to the left unit caused lateral excitatory and inhibitory currents to enter the center unit. These currents to the center unit were designed to be similar to the currents entering the single-unit in response to auditory stimuli. We performed the same procedure for auditory inputs to the right unit: lateral excitatory and inhibitory currents from the right were designed to enter the center unit in a manner similar to the single-unit case. Using this procedure, we extended the differential SST and PV inhibition in the single-unit model to work in the case of a tonotopy without the need for exhaustive parameter fitting.

The oddball stimulus activated the neighboring units to the center one, whose activity we measured (Stimulus described in Table 2, Figure 4A). We observe thalamic depression in the thalamus which decreases responses to the repeated activation from one unit, and large responses to the unrepeated tone. We simulated PV and SST suppression with the functions *I*_Opt,PV_(*t*) and *I*_Opt,SST_(*t*). These functions were turned on 100ms before tone onset and turned off 100ms after tone onset. In the rate model, the inhibition was dimensionless and free parameters chosen to be *I*_Opt,PV_(*t*) = −4 during PV inhibition and *I*_Opt,SST_(*t*) = −2 during SST inhibition. In the spiking model, we chose *I*_Opt,PV_(*t*) = −0.2nA during PV inhibition and *I*_Opt,SST_(*t*) = −1nA during SST inhibition (Figure 4B—G).

**Figure 4.**
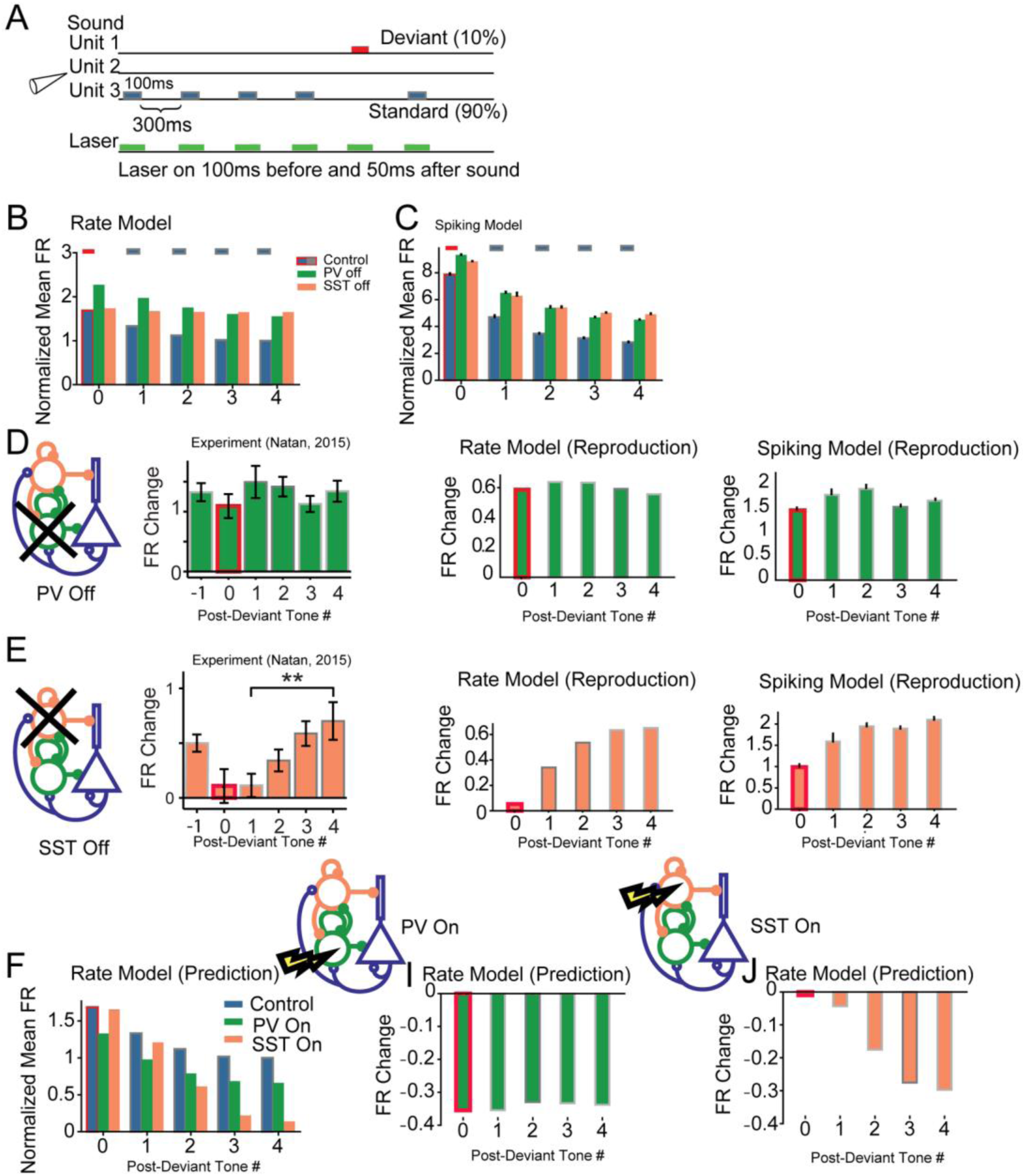
Summary of SSA in the rate and spiking model. A: Oddball stimulus consisted of two tones: standard tones (gray) appear with 90% probability, whereas deviant tones (red) appear with 10% probability. B, C: Average response of the excitatory population to the deviant (red outline) and subsequent standards (gray outline) without stimulation; with PV suppression (green) and with SST suppression (orange). B. Rate model. C. Spiking model. D. Additive change in response of excitatory population due to PV suppression in the rate and spiking models to the deviant (red outline) and standards (gray outline). Left: from published data. Center: rate model. Right: Spiking model. E. Additive change in response of excitatory population due to SST suppression in the rate and spiking models to the deviant (red outline) and standards (gray outline). Left: from published data. Center: rate model. Right: Spiking model. F. Predictions for the responses to the oddball stimulus with and without interneuron activation in the rate model. Left: Mean responses of the excitatory population to the deviant and subsequent standards (red/gray outline: no activation; green: PV activation; orange: SST activation). Middle: Change in excitatory neuron responses due to PV activation, Right: Identical plot as the middle panel, but for SST activation. PV activation resulted in a near-uniform decrease in FRs, whereas SST resulted in an increase in adaptation.

In the rate and spiking model, the firing rates increased uniformly across all post-deviant tones (Figure 4D,E). In the rate and spiking model, the firing rates exhibited an increase in disinhibition as a function of post-deviant tone number (Figure 4F,G). Both results agree with existing results in SSA (Natan et al. 2015). In order to establish the robustness of these results, we varied several parameters and measured the Common-contrast SSA Index (CSI) (Yarden and Nelken 2017),

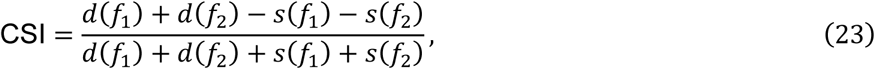

where *d*(*f*_*i*_) is the deviant rate response and *s*(*f*_*i*_)) is the standard rate response to frequency *f*_*i*_. For full adaptation, when the standard responses are 0, CSI = 1, indicating a high degree of SSA. If the standard responses are equal to the deviant responses, then CSI = 0, indicating a low degree of SSA.

We performed a parameter sweep with four key parameters of circuit connectivity (Figure 5): (1) recurrent excitation (*w*_*ee*_, Figure 5A,B,E,F), a key parameter considered in many studies (Yarden and Nelken 2017), especially those related to inhibitory stabilized networks (ISNs) (Tsodyks et al. 1997; Litwin-Kumar, Rosenbaum, and Doiron 2016); (2) timescale of thalamic depression (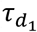, Figure 5C,D,G,H) whose reported values vary over a large range, from 0.8s to 3s (Yarden and Nelken 2017; Natan et al. 2015); (3) the strength of PV activation or inactivation (Figure 5A,C,E,G); and (4) the strength of SST activation or inactivation (Figure 5B,D,F,H). In all cases, inactivating SSTs reduced the CSI relative to PV inactivation, reflecting the increasing disinhibition over post-deviant tones. The x-axis of each plot corresponds to the strength of the optogenetic laser. Negative values correspond to a decrease in inhibitory activity, and positive values correspond to an increase in inhibitory activity. These effects may be more clearly seen in the model equations (Equations 1, 5, and 13), where we add the terms *I*_Opt,PV_(*t*) and *I*_Opt,SST_(*t*).

**Figure 5.**
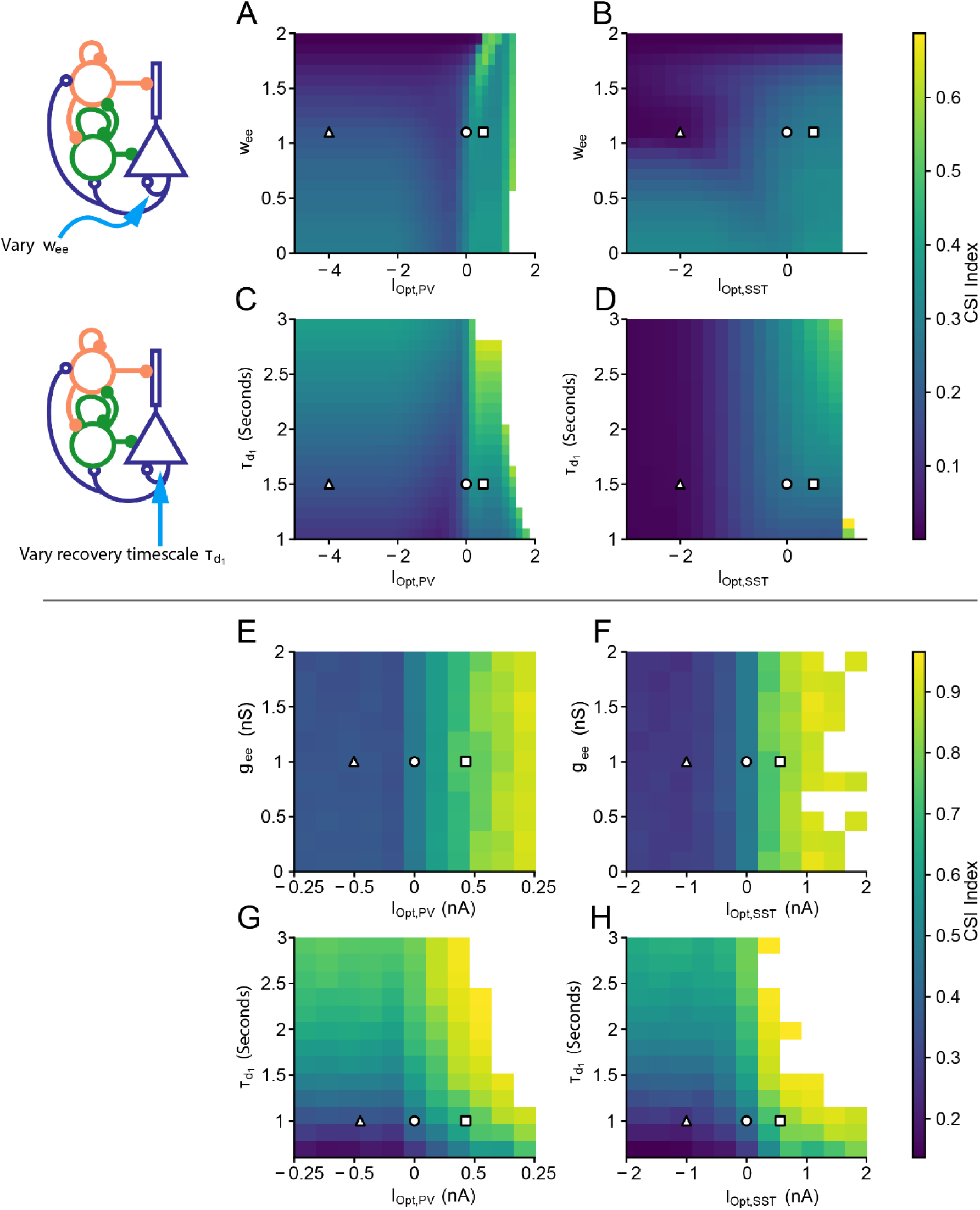
Predicted effects of the key parameters on SSA index (CSI) for the rate model (A-D) and spiking model (E-H). Control parameters are denoted by white circles, PV and SST inactivation parameters are denoted by white triangles, and PV and SST activation parameters are denoted by white squares. The control parameter values, I_Opt,PV_ = I_Opt,SST_ = 0, are denoted by white circles. A, E: PV optogenetic parameter vs recurrent excitation (w_ee_). B, F: SST optogenetic parameter vs recurrent excitation. C, G: PV optogenetic parameter vs thalamic depression time constant 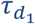. D, H: SST optogenetic parameter vs thalamic depression time constant. White regions in all subfigures denote areas where the firing rate (FR) of the standard tone is too low (FR< 0.1), or where the excitatory response saturates, making CSI measurements impossible.

This analysis reveals robustness in parameter ranges for given optogenetic modulation strengths. The CSI in the control case (white circle) changed little when the parameters *w*_*ee*_ and 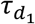 were varied for a given optogenetic strength. In other words, for a fixed value on the x-axis, changing positions in the vertical direction on each plot did not change the CSI index significantly for a nontrivial range of parameter values. This observation suggests that the cortical model can operate in a broad parameter regime, and precise parameter values may not be important for normal function. In extreme cases, decreasing recurrent excitation removed the decrease in CSI following SST inactivation (Figure 5B), suggesting that sufficient recurrent excitation is an important factor in generating responses in the SSA paradigm. Second, while increasing optogenetic inhibition had little effect on the CSI, increasing optogenetic activation showed an increase in CSI in all cases (CSI= 0.35 for PV activation and CSI= 0.31 for SST activation). Therefore, we predicted that optogenetic activation of PVs and SSTs will generally improve context-dependent cortical responses.

Like the rate model, the spiking model exhibited little sensitivity to changes in *w*_*ee*_ and 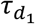. However, the spiking model showed almost no dependence on recurrent excitation *w*_*ee*_ in the case of SST inactivation (Figure 5E). This effect is likely due to the differences in connectivity between the rate and spiking models. In the rate model, lateral connections depend entirely on excitatory activity, thus SSA results in the rate model are more sensitive to changes in recurrent excitation. In the spiking model, recurrent excitation plays a less important role because the lateral connection probabilities are low (*p* = 0.1), whereas the connection probabilities within units are high (*p* = 0.6).

The three-unit model was developed to reproduce the compensating mechanisms of the single-unit model: PV suppression results in constant disinhibition for repeated tones, and SST suppression results in a compensating effect from PVs before adaptation that weakens as adaptation strengthens. These differential roles explain experimental data in the SSA paradigm to a remarkable degree. We then asked whether this simple mechanism is sufficient to reproduce additional optogenetic experiments. For the remainder of the paper, we use the three-unit rate model with no parameter modifications except for the changes in the inhibition modes and the auditory inputs that depend on the experimental paradigm (Table 2, Figure 1C, and Equation 10).

### Differential effects of inhibitory neuron manipulation on cortical forward suppression

Context dependence of auditory responses has been revealed on many time scales. In a well-studies phenomenon termed “forward suppression”, the responses of AC neurons to a tone are suppressed if the tone is preceded by another tone, but the level of suppression depends on the frequency difference between the two tones (Figure 6). In the experiment, the first tone, called the masker, varies in frequency between trials, while the second tone, called the probe, remains fixed at the preferred frequency of the neuron. This phenomenon was explained by feedforward depression, but the inhibitory neurons were recently shown to also control forward suppression (Phillips, Schreiner, and Hasenstaub 2017). PV inactivation (orange) concurrent with the auditory stimulus resulted in a selective increase in forward suppression at the preferred frequency relative to the control case (blue), whereas SST inactivation (green) reduced forward suppression at the preferred frequency relative to the control case (blue) (Figure 4B second row) (Phillips, Schreiner, and Hasenstaub 2017).

**Figure 6.**
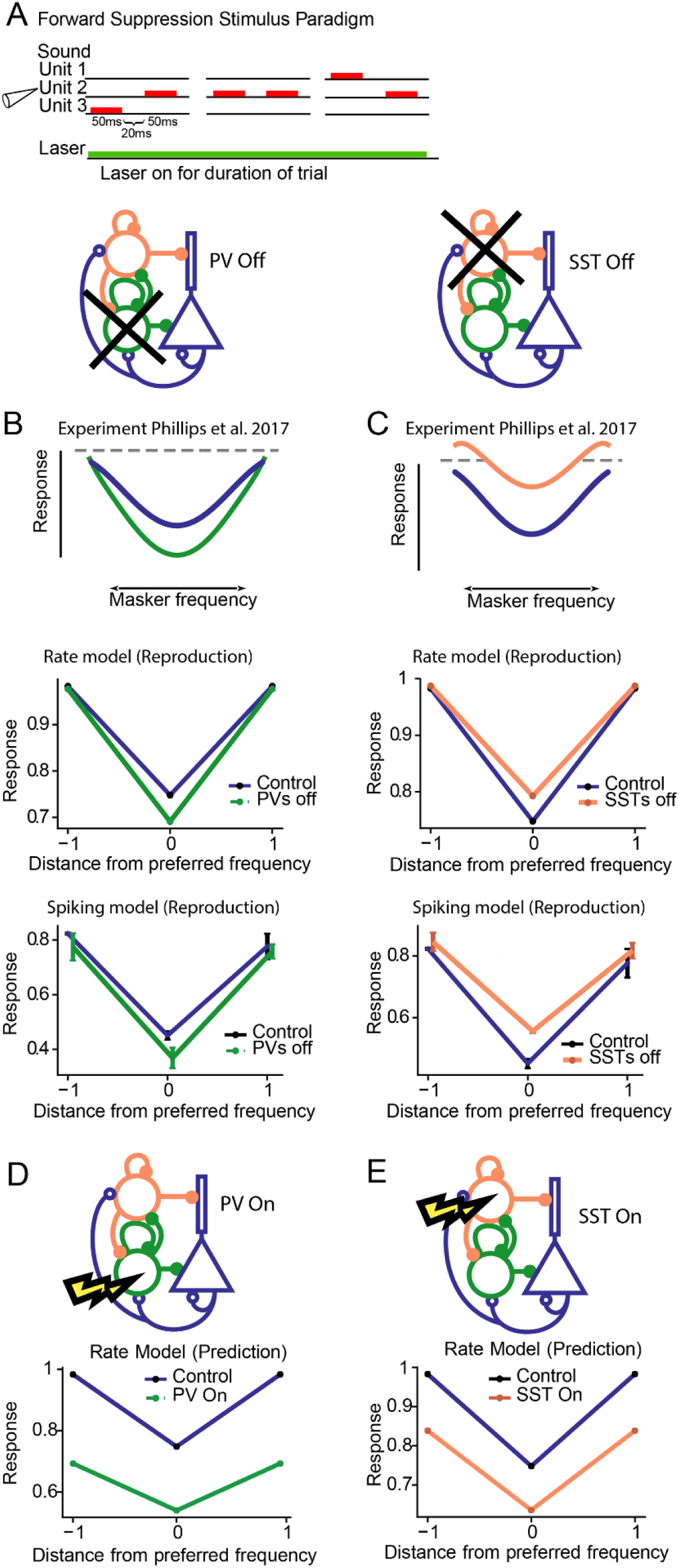
Forward suppression in the rate and spiking model. A. The stimulus consisted of pairs of tones activating either neighboring or the same iso-frequency units. The laser was presented continuously throughout stimulation trials. B,C,D, E: Responses of excitatory neurons to the probe tone as a function of the frequency of the masker. Blue: control. B, C: Top row: Schematic of results from Phillips et al., 2017. Middle row: Results from the rate model. Bottom row: Results from the spiking model. B: Results of PV suppression (green). C: Results of SST suppression (orange). D,E: Rate model prediction for forward suppression during PV and SST activation. PV and SST activation resulted in enhanced forward suppression. D: Results from PV activation. E. Results from SST activation.

We used the same parameters for connectivity within the circuit as with SSA to reproduce the experimental findings, with only slight changes to the input strength (*q* = 1.3). The stimuli used in the forward suppression paradigm place the baseline state in the strong inhibitory regime (Figure 1C). Both the rate (Figure 6A middle, 6B middle) and spiking models (Figure 6B bottom 6B bottom) yielded the existence of experimentally observed differential effects for PV (Figure 6A) and SST inactivation (Figure 6B): PV inactivation drove a selective decrease in responses whereas SST inactivation drove a suppression of excitatory neuronal responses. We do not expect the scales between the rate and spiking models to match precisely, and we only sought to match the existence of the differential phenomena. Functions *I*_Opt,PV_(*t*) and *I*_Opt,SST_(*t*) were turned on for the duration of the simulation to match the experimental protocol. In the rate model, the inhibition was dimensionless and free parameters chosen to be *I*_Opt,PV_(*t*) = −4 during PV inhibition, *I*_Opt,PV_(*t*) = 0.5 during PV activation, *I*_Opt,SST_(*t*) = −2 during SST inhibition, and *I*_Opt,SST_(*t*) = 1.2 during SST activation. In the spiking model, we chose *I*_Opt,PV_(*t*) = −0.2nA during PV suppression, *I*_Opt,PV_(*t*) = during PV activation, *I*_Opt,SST_(*t*) = −1nA during SST inhibition, and *I*_Opt,SST_(*t*) = −1nA during SST activation.

At first glance, this result seems paradoxical given that PV suppression generally results in excitatory disinhibition as shown in the adaptation and SSA results (Figures 3,4), but the underlying mechanism is straightforward to understand. Following PV suppression, excitatory activity is indeed disinhibited, but the firing rate function of the excitatory population saturates. This behavior means that higher activity neurons are generally disinhibited less strongly than lower activity neurons. Thus, upon receiving the second tone, the input received by the excitatory population is weaker due to thalamic depression, but the disinhibition is greater relative to the disinhibition in the first tone. This phenomenon also requires a special property in the thalamic depression variable *g*(*t*), namely that it cannot depress too quickly during the first time or else the excitation in the second tone will be too weak, and although it recovers slowly, it recovers enough such that the input from the second tone is still somewhat strong. In the case of SST suppression, PVs compensate for the loss of inhibition in the first tone, but lose the ability for compensation in the second tone, so Exc are able to respond more strongly relative to the control case. Thus, forward suppression is weakened.

Next, we tested the effects of activating PVs or SSTs (as could be done with ChR2 experimentally) on model responses. The model predicted that both PV and SST activation will result in an increase of forward suppression across preferred and sideband frequencies (Figure 6D,E).

### Differential adaptation to repeated tones along the frequency response function

Neurons in A1 adapt to repeated tones (Natan et al. 2015). This adaptation is proportional to the strength of their tone-evoked responses: it is stronger in the center of the frequency response function, and weaker for the sidebands (Natan, Rao, and Geffen 2017). A recent study found that PVs and SSTs exert a differential effect on this form of adaptation: Suppressing PVs drives disinhibition selective to the sidebands in the adapted state, whereas suppressing SSTs drives disinhibition both in the center and at the sidebands of the frequency response function of excitatory neurons (Natan, Rao, and Geffen 2017). To understand how inhibitory neurons affect adaptation across different frequency-tuned inputs, we presented a sequence of 8 tones at each frequency to generate adapting tuning curves (Figure 7A), and repeated this stimulus with PV and SST suppression for the model circuit. We found that this auditory paradigm resulted in a below-threshold integration of 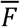, so the system switched to a state of strong baseline inhibition (and importantly, the model did not respond in precisely the same way as in SSA and forward suppression).

**Figure 7.**
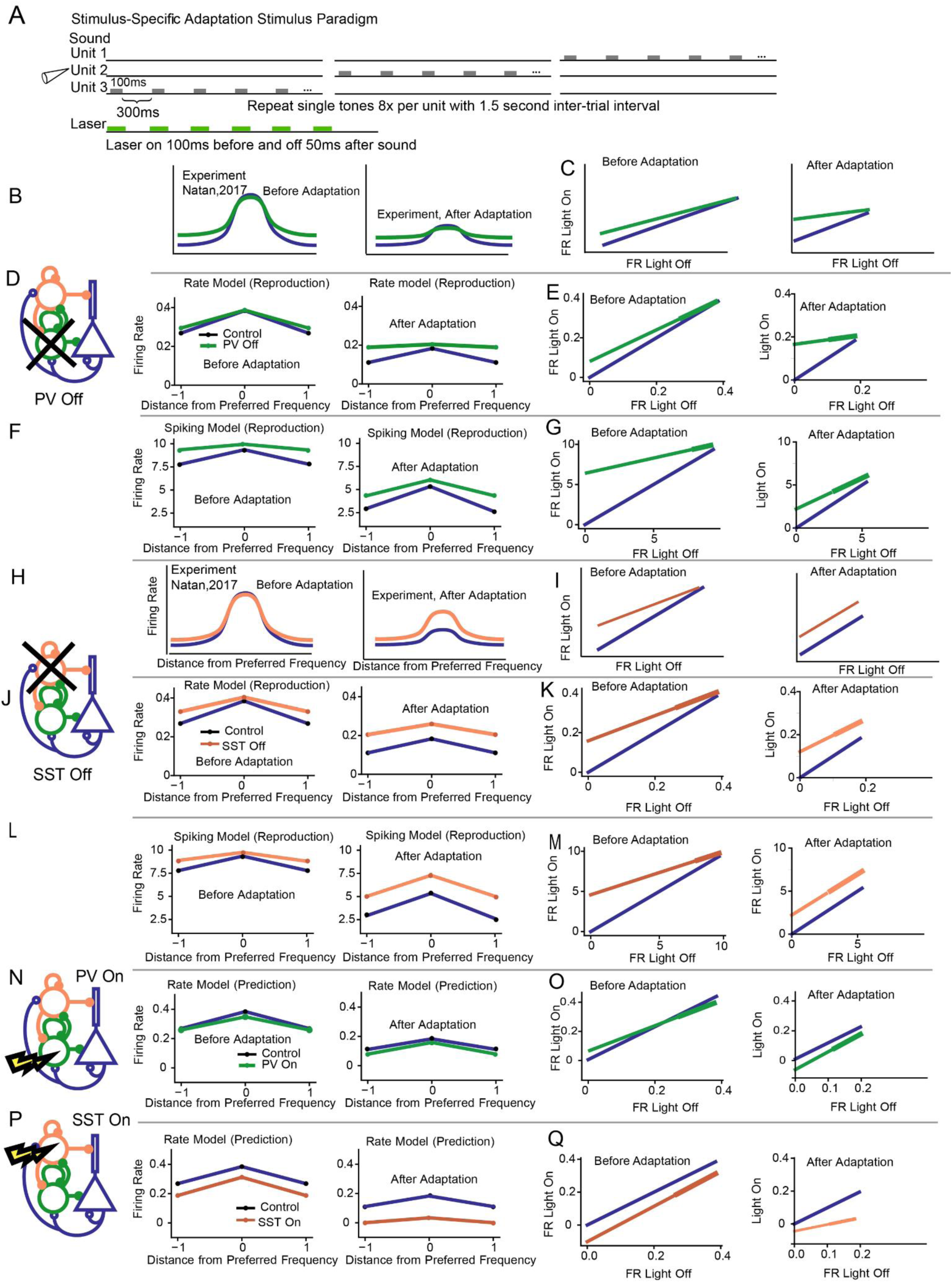
Adaptation to repeated tones along the frequency response function. A. The stimulus consisted of a sequence of repeated tones, presented to each iso-frequency unit. On stimulation trials, the laser overlapped with the sound stimulus. B, D, F, H, J, L, N, P: The responses of excitatory units to the first (left) and last tone (right) as a function of the distance in frequency between the unit and the stimulus without (blue) and with (green: PV suppression; orange: SST suppression) stimulation. C, E, G, I, K, M, O, Q: The response of excitatory neurons to tone 1 (left) and tone 8 (right) on light on and light off trials. The control lines have unit slope because light on and light off in an experimental condition yields no changes to the firing rate. Thicker lines in Figure 7E,G,K,M represent the peak excitatory responses from the first and last simulations taken directly from the simulations, whereas thinner lines are linear extrapolations to assist the visual comparison to the control line (blue). B,C. Experimental results, PV suppression. D,E: Rate Model, PV suppression. F,G. Spiking model, PV suppression. H,I. Experimental results, SST suppression. J,K. Rate model, SST suppression. L,M. Spiking model, SST suppression. N,O. Rate model, PV activation. P,Q. Rate model, SST activation.

Our model reproduced the differential experimental effects of PV and SST suppression (Figure 7). In the rate model before adaptation, PV and SST inactivation resulted in sideband disinhibition with little to no disinhibition at the preferred frequency (Figure 7D,J eft). After adaptation, PV inactivation resulted in sideband disinhibition and no preferred frequency disinhibition (Figure 7D right), whereas SST inactivation resulted in disinhibition across all sideband and preferred frequencies (Figure 7J right). The spiking model closely mirrored these results (Figure 7F,L). The ratio of excitatory responses between light off and light on trials summarize the degree of sideband and preferred frequency disinhibition (Figure 7E,G,K,M).

The mechanisms behind these results involve synaptic facilitation and depression and the compensating mechanism discussed in the earlier sections for SSA and adaptation. In the case of PV suppression, SSTs were the only interneurons capable of contributing to Exc inhibition, so only Exc-SST interactions drove the observed effects. In particular, lateral SST to Exc synapses suppressed the center unit over each tone, and facilitation allowed this suppression to persist throughout adaptation. Note that this preferred-frequency effect was not observed in SSA because we never directly stimulated the center unit. Next, in the case of SST suppression, the increasing disinhibition with adaptation at the preferred frequency was a consequence of the same compensating mechanism as in SSA. The tuning curves shown in Figure 7B,H are representative cartoons of the changes in the tuning curves observed quantitatively (Natan, Rao, and Geffen 2017), but we aimed to reproduce the light-on and light-off curves (Figure C,H) using our model. The comparison of the model tuning curves to the cartoons from the experiments appear to be significantly different at the sidebands, but in fact our model outputs closely match the desired lines in Figure 7C,I.

Turning to predictions, we found that before adaptation, PV activation resulted in a slight decrease at the preferred frequency, whereas SST inactivation reduced overall firing rates across all frequencies (Figure 7N,O). After adaptation, PV and SST activation resulted in a subtractive effect. General optogenetic activation and inactivation of PVs and SSTs modulated tuning-curves in combinations of additive, subtractive, multiplicative, and divisive effects (Phillips and Hasenstaub 2016). Our model reproduced one of the key results in these studies as well, since PV and SST inactivation were found to have additive and divisive effects on the frequency response functions of excitatory neurons (Figure E,G,K,M,O,Q).

Similar to SSA, the functions *I*_Opt,PV_(*t*) and *I*_Opt,SST_(*t*) were turned on 100ms before tone onset and turned off 100ms after tone onset. In the rate model, the inhibition was dimensionless and free parameters chosen to be *I*_Opt,PV_(*t*) = −0.5 during PV inhibition, *I*_Opt,PV_(*t*) = 1.2 during PV activation, *I*_Opt,SST_(*t*) = −1 during SST inhibition, and *I*_Opt,SST_(*t*) = 0.1 during SST activation. In the spiking model, we chose *I*_Opt,PV_(*t*) = −1nA during PV inhibition and *I*_Opt,SST_(*t*) = −1nA during SST inhibition.

### PVs Enhance Feedforward Functional Connectivity

Cortical neurons in AC receive inputs from the thalamic auditory nuclei. As the result, neuronal responses in the cortex are correlated with neuronal firing in the thalamus. These interactions can be captured using an Ising model to measure the connection from the thalamus to the cortex. When PVs were activated, the functional coupling between cortical and thalamic responses (Hamilton et al. 2013) became stronger. The specific mechanism underlying this change is unknown.

Using the three-unit model, we identified a candidate mechanism for the enhanced thalamo-cortical correlation following PV activation. We assumed that the feedforward functional connection from the thalamus to the cortex is the same as the anatomical connection, so thalamic inputs directly modulated cortical responses in our model. Following an increase in inhibition, cortical responses became sharper, thus aligning more closely with thalamic inputs and improving feedforward functional connectivity (Figure 8).

**Figure 8.**
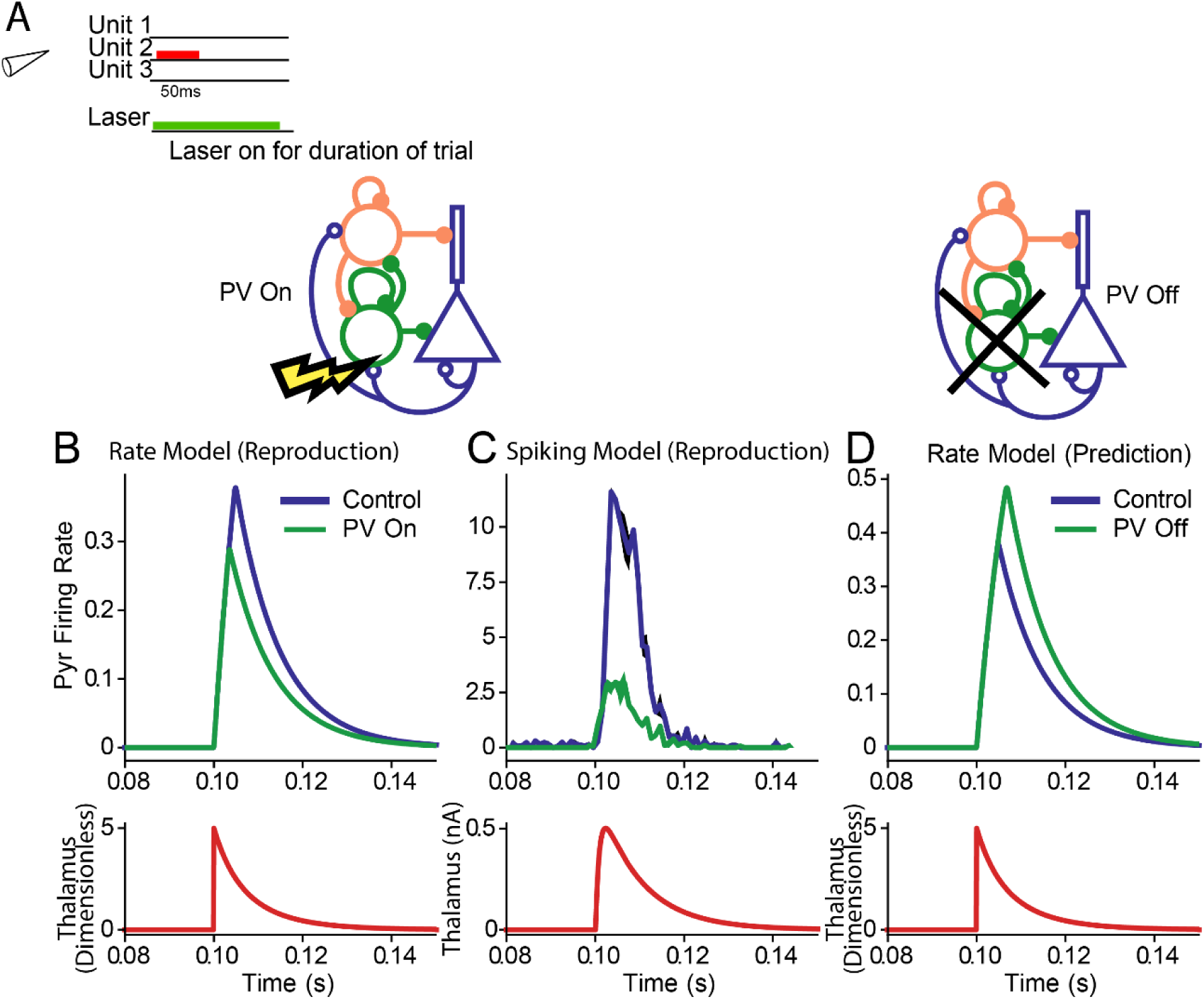
Activation of PVs enhanced feedforward connectivity in the model. A. Stimulus was a single tone accompanied by a laser on stimulation trials presented at 0.1 s. B. Top: Cortical excitatory population responses to tones without (blue) and with PV stimulation (green) in the rate model. Bottom: Thalamic input (red). C. Top: Cortical excitatory population responses to tones without (blue) and with PV stimulation (green) in the spiking model. D: Rate model prediction for effects of PV inactivation.

PV activation (green) in the rate model resulted in an increase in the Pearson correlation between the control (blue) and thalamic inputs (red), from 0.77 and 0.83 (Figure 8B). Thus, whereas inhibitory activation decreased the overall firing rate, the response became more synchronized to the thalamic inputs, resulting in an increase in feedforward functional connectivity. In the spiking model, PV activation resulted in a delayed response of excitatory activity, but we were interested in tested whether PV-activated Exc response profile resembled the thalamic activity more than the control Exc response. To make this comparison, we shifted the PV trace so that the onset of PV-activated Exc activity (green) coincided with the onset of the control curve (blue) (Figure 8C. An equivalent approach would be to measure the peak value of the cross-correlation between excitatory and thalamic activity without shifting the data in time). We observed an increase in the Pearson correlation from 0.87 in the control Exc activity to 0.95 in the PV-activated Exc activity, thus demonstrating a sharpening of excitatory responses, and an increase in feedforward functional connectivity. As in the previous paradigms, we only aimed to show the existence of changes without requiring the magnitude of change to match the experimental data.

These results provide for a simple plausible mechanism for enhanced feedforward functional connectivity: as inhibition reduces the overall cortical inputs, cortical responses better synchronize to thalamic inputs, resulting in stronger correlated activity. We remark that the classic approach to the problem of establishing proper functional connectivity relies on sophisticated models such as Ising models (Ganmor, Segev, and Schneidman 2011), expectation-maximization (Turaga et al. 2013), and subspace identification (Nonnenmacher, Turaga, and Macke 2017), because direct calculations of correlations can lead to false positives when the anatomical connections are not known (Hamilton et al. 2013). Our model has explicit anatomical connections, which eliminates the problem of false-positive correlations. Thus, in this case, the use of the correlation serves as a reliable proxy for the Ising model.

### Balanced Networks

Multiple studies postulated that excitatory and inhibitory currents are matched in cortical circuits, contributing to stability of circuit function (Denève and Machens 2016; Zhou and Yu 2018). Therefore, we tested whether our model operated as a balanced network. To make this measurement, we took ratios of inhibitory and excitatory currents during suprathreshold activity, because our model ignores virtually all subthreshold activity. Interestingly, with no tuning, our models showed evidence of operating as a balanced network: as we varied the input strength to the rate and spiking models (Figure 9C,F), the ratio of excitatory to inhibitory inputs to the excitatory population (Figure 9B,E) remained constant. The rate model had an excitatory/inhibitory ratio of 0.37 (Figure 9B), and the spiking model had an excitatory/inhibitory ratio of 2.5 (Figure 9E). These results suggest that excitatory-inhibitory balance a robust, emergent feature of cortical networks. The large differences in scales are to be expected when comparing a dimensional and dimensionless model.

**Figure 9.**
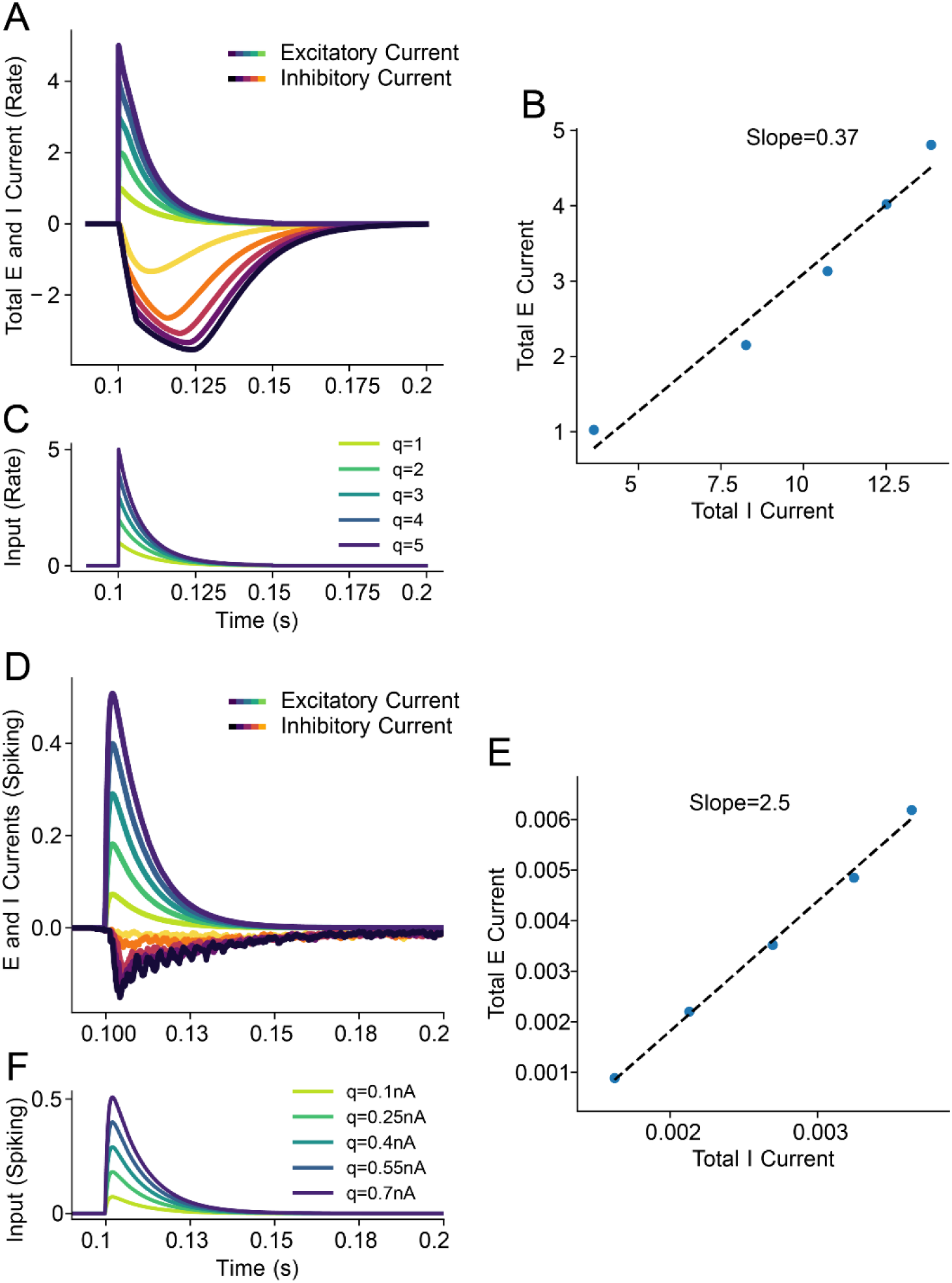
Excitatory-inhibitory balance in the rate and spiking models A. Plot of incoming excitatory and inhibitory currents into the Exc population as a function of different input strengths (C). Darker currents correspond to stronger inputs B. A best-fit line (dashed) accurately captures the ratio of excitatory and inhibitory responses, implying excitatory-inhibitory balance. Equivalent results for the spiking model are shown in panels D, E, and F.

## Discussion

A wealth of recent studies provide evidence for distinct function of different types of cortical inhibitory neurons in temporal processing of auditory information. The studies demonstrate that different types of inhibitory neurons, SSTs and PVs, play a differential role in auditory processing, controlling adaptation at different time scales and contexts, and changes to feedforward functional connectivity. Our goal was to integrate the results of these studies to understand whether the observed effects were due to a small set of mechanisms.

We built an idealized rate and spiking model that reproduced multiple key results from studies that tested the function of specific inhibitory opsins in specific cells in the auditory cortex. In addition to including different baseline states that modulate the strength of PV-to-Exc and SST-to-Exc synapses, the key mechanisms underlying our models included the fast temporal activation of PVs, the delayed, broad temporal activation of SSTs, the ability for PVs to compensate for weakened SST activity, and dynamic synapses including SST-to-Exc facilitation. These interactions accounted for the differential modulation of cortical responses by interneuron subtypes and suggests that a simplified set of mechanisms can support experimental results.

To reproduce the differential function of SSTs and PVs in stimulus-specific adaptation, we built a model loosely based on multiple existing models for SSA and multiple configurations of spiking neuron populations, consisting of inhibitory and excitatory neurons. Previously, a two-layer rate model with synaptic depression was proposed to establish the relationship between the cortical response and the parameters in SSA experiments, such as stimulus frequency differences, probability of deviation, and tone presentation rate. Yarden et al. (2017) successfully used a multi-unit rate model arranged in a coarse tonotopy consisting of inhibitory and excitatory populations to reproduce general deviance detection, but model has not yet been adapted to explain differential interneuron modulation. Another existing model of SSA including differential inhibitory modulation demonstrating similar differential inhibitory effects as in our SSA result (Figure 4), but did not include a tonotopy (Natan et al. 2015). These models only included one type of inhibitory neuron type or did not include tonotopy, and therefore could not account for the observed differential effects of suppression of SSTs and PVs on SSA across multiple frequencies. In the present study, we developed a simple rate and spiking model that accounted for multiple inhibitory cell types and which faithfully reproduced the differential effects of SST and PV inactivation in SSA (Figure 4). In addition, a parameter sweep revealed that both the rate and spiking models were robust to large changes in key parameters commonly explored in the literature, suggesting that SSA is a robust phenomenon (Yarden and Nelken 2017).

Existing models that reproduce the enhanced forward suppression from PV inactivation and the reduced forward suppression from SST inactivation (Figure 6) include multiple layers that require both depression and facilitation (Phillips, Schreiner, and Hasenstaub 2017), or rely on depressing recurrent excitation and do not distinguish between inhibitory subtypes (Loebel, Nelken, and Tsodyks 2007). We incorporated depression and facilitation in the model synapses and reproduced the former results with only a single layer, suggesting a surprisingly simple mechanism supporting forward suppression. Furthermore, the models in the present study reproduced tuning-curve adaptation effects previously observed experimentally but not computationally: SSTs exhibited strong preferred-frequency disinhibition following adaptation, while PV disinhibition is independent of the degree of adaptation (Figure 7) (Natan, Rao, and Geffen 2017). These results suggest that the underlying mechanism(s) of the model, namely the PV/SST compensation effect, combined with the facilitating SST-to-Exc synapse, may serve as a general mechanism of adaptation. In addition, our models reproduced changes in feedforward functional connectivity (Figure 8). By increasing PV activity in the models, excitatory activity decreased but became more time-locked to thalamic inputs. This effect agreed with observations in the cortex, where PV activation resulted in enhanced feedforward functional connectivity (Hamilton et al. 2013). The effects of inhibition on sharpening cortical responses have been well-established, thus our models serve as plausible mechanisms for this change (Wehr and Zador 2003; Cardin et al. 2009; Sohal et al. 2009). Finally, our models were shown to operate as a balanced network, where inhibitory and excitatory currents entering neural populations were shown to maintain a consistent ratio across input strengths, suggesting a generality to the theory of balanced networks.

One drawback of the model is that it does not feature population spikes, which explain many fundamental cortical responses in AC (Wehr and Zador 2003). In future work, we will seek to reconcile the differences between our models and the population spike model of SSA (Loebel, Nelken, and Tsodyks 2007). Establishing the importance of depression and facilitation in different synapses and extending our model to include population spikes warrants further study.

Although we do not explore simultaneous auditory stimuli in this study, it is worth mentioning the response properties of the network due to recent interest in supralinear network models (Rubin et al. 2016). Throughout this paper, neurons operate in a linear manner when above threshold: neurons add inputs linearly, until the maximum rate is reached in the rate models, or until the refractory period saturates spiking rates in the spiking model. The models do not use sub-threshold responses to modulate population activity.

Multiple studies from different laboratories revealed the differential effect of distinct inhibitory neurons in auditory processing. We show that a minimalistic model, built on simple mechanisms, is capable of reproducing disparate results in the literature. As inhibitory neurons form similar circuits throughout the mammalian cortex, this model can be readily adapted to test their function and generate predictions (with adjustments for local changes in connectivity) in different sensory modalities.

## Acknowledgments

This work was supported by National Institutes of Health (NIH R01DC014479, NIH R01DC015527), Human Frontier in Science Foundation Young Investigator Award and the Pennsylvania Lions Club Hearing Research Fellowship to MGN. The authors thank the members of the Geffen laboratory and the Hearing Research Center at the University of Pennsylvania, Jason Kim and Lia Papadopoulos, for insightful comments on the manuscript and discussions.

